# Assessing Scale and Predictive Diversity in Models for Single-Cell Transcriptomics based on Geneformer

**DOI:** 10.1101/2025.11.04.686458

**Authors:** Junfan Chen, Fabian Schmidt, Ricardo Henao

**Affiliations:** Biological and Environmental Science and Engineering Division, King Abdullah University of Science and Technology (KAUST), Thuwal, Makkah, Saudi Arabia; Department of Biostatistics & Bioinformatics, Duke University, Durham, North Carolina, United States

## Abstract

Foundation models are increasingly applied to single-cell transcriptomics, where they promise to capture generalizable representations that support diverse downstream analyzes. However, two central questions remain: Does scaling pre-training data reliably improve performance, and do models trained on rank-ordered expression profiles confer advantages for mitigating batch effects? We addressed these questions by systematically assessing transformer-based models pre-trained on ranked single-cell profiles with varying data scales. The models were evaluated on masked gene prediction and downstream tasks, including cell type classification, perturbation response prediction, and zero-shot batch integration. To complement prediction accuracy, we further quantified prediction repetition, uniqueness, and diversity. In addition, we evaluated architectural refinements that incorporate cumulative prediction adjustment and similarity-based regularization; however, the study mainly focused on comparative evaluations rather than the development of new models. Our results indicated that scaling pretraining corpora improved masked prediction accuracy but did not consistently enhance downstream performance. Smaller models often matched or exceeded larger ones, indicating diminishing returns relative to scale. Rank-based models offered limited robustness to batch effects and consistently underperformed relative to domain-specific correction methods. Across all scales, high redundancy in predicted genes remains a major limitation. Together, these findings challenge assumptions that larger datasets or rank-order modeling automatically confer stronger generalization. Progress in single-cell foundation models may depend less on scale and more on pre-training objectives that enhance predictive diversity and biological plausibility.

**Author summary:** In recent years, large machine learning models have been adapted to study single cells, raising hopes that simply training on more data would automatically lead to better biological insights. Our study aimed to test this assumption by carefully evaluating models trained on small versus very large collections of single-cell data. We also asked whether models that learn from ranked lists of gene expressions offer an advantage in correcting technical differences between experiments, a common challenge in biology.

We found that bigger is not always better. Although larger models improved performance in predicting missing genes, this advantage did not consistently translate to stronger results in tasks such as cell type classification or data integration across studies. In fact, smaller models often matched or outperformed their larger counterparts. Models trained on ranked data were also not sufficient to reliably correct batch effects, and traditional tools still worked better for this purpose.

Our findings suggest that future progress may come less from ever-larger models and more from rethinking how models learn from biological data. This work provides guidance for researchers designing new approaches and helps clarify realistic expectations for applying artificial intelligence to single-cell biology.

## Introduction

In recent years, foundation models, large-scale pre-trained models capable of learning universal feature representations and transferable across downstream tasks, have become a dominant paradigm across multiple fields [1–3]. Single-cell transcriptomic data [4, 5], with its high dimensionality, complexity, and scale, provides an ideal substrate for such approaches. Advances in single-cell sequencing have enabled the systematic characterization of cell states and functions at single-cell resolution; yet traditional bioinformatics methods often struggle with efficiency and scalability when applied to increasingly large datasets. Foundation models offer a promising alternative, as demonstrated by representative efforts such as Geneformer (GF) [6], scBERT [7], scGPT [8], and scFoundation [9], which are trained on tens of millions of single-cell profiles and show strong performance in tasks including cell type classification and gene expression prediction.

Among these, GF was the first to adapt the Bidirectional Encoder Representations from Transformers (BERT) architecture [10] to rank-ordered single-cell transcriptomic data. Early studies highlighted its transfer learning ability and suggested its potential to alleviate batch effects. However, subsequent evaluation efforts have revealed limitations. For perturbation response prediction, its performance is comparable to, or worse than lightweight models such as Multilayer Perceptrons (MLPs) or GEARS [11, 12]. In cell type annotation and marker gene identification, it has not consistently outperformed traditional methods such as Differential Expression Tests (DET) or scANVI [13–15]. In zero-shot batch correction, it falls markedly behind specialized approaches such as scVI [15–17]. Insights from natural language processing further suggest that scaling pre-training data alone does not guarantee stronger models and may even impair generalization [18–20]. Moreover, GF’s masked language modeling strategy [10] introduces biological inconsistencies; namely it often predicts the same gene repeatedly within a single profile and shows a bias toward high frequency and ubiquitously expressed genes, reducing prediction diversity and limiting biological interpretability.

This study focuses on GF and investigates two central questions: (1) Does scaling pre-training data consistently improve downstream performance on single-cell transcriptomic data? and (2) Do models trained on rank-ordered profiles truly confer advantages in mitigating batch effects? To address the issue of biological inconsistency, we propose structural refinements to the GF framework, hereafter referred to as GF_CAB_, Geneformer with **C**umulative-assignment on the prediction **A**djustment and **B**oosting on the similarity-based diversity, and systematically evaluate their impact across a range of tasks.

Our experiments show that increasing the size of the dataset used for pre-training yields only marginal gains in certain classification tasks and can even degrade performance in batch-effect correction. In contrast, GF_CAB_ mitigates several shortcomings of the original approach and shows reduced dependence on large-scale pretraining: it achieves performance comparable to GF trained on datasets over an order of magnitude larger and demonstrates modest improvements in batch effect mitigation. However, it still trails behind task-specific bioinformatics methods. Together, these findings reinforce the view that “more pre-training data does not necessarily produce stronger models” in the biological context and highlight the limitations of rank-ordered foundation models in terms of biological consistency and batch handling. Although our refinements do not yet surpass traditional tools, their effectiveness on smaller datasets provides new empirical evidence and a foundation for further optimization of single-cell foundation models.

## Results

We evaluated both the original GF and our refined GF_CAB_ models on pre-training objectives and a diverse set of downstream tasks. These tasks included three classification benchmarks (token-wise gene dosage sensitivity, tissue-specific cell type, and cardiomyocyte subtype prediction) along with zero-shot batch-effect evaluation across four independent single-cell datasets (Immune 330K, Pancreas 16K, PBMC 12K and the scCello cell type dataset [15]). This evaluation revealed the relative contributions of structural refinements to predictive diversity and batch-effect robustness, providing a systematic comparison of rank-ordered foundation models in biologically realistic settings.

### Pretraining performance

We first assessed the models on their pretraining objectives by quantifying three aspects of masked token prediction: accuracy, repetition, and uniqueness (Fig 2B–D, with detailed scores in S1 Table). Accuracy measured the proportion of correctly recovered masked genes, repetition captured redundancy in predictions across masked positions or overlap with unmasked input genes, and uniqueness reflected the proportion of distinct genes predicted, providing a proxy for diversity. To contextualize these metrics, we also recorded relative training times (Fig 2A), thereby evaluating the trade-offs between predictive performance and computational demand.

**Fig 1.**
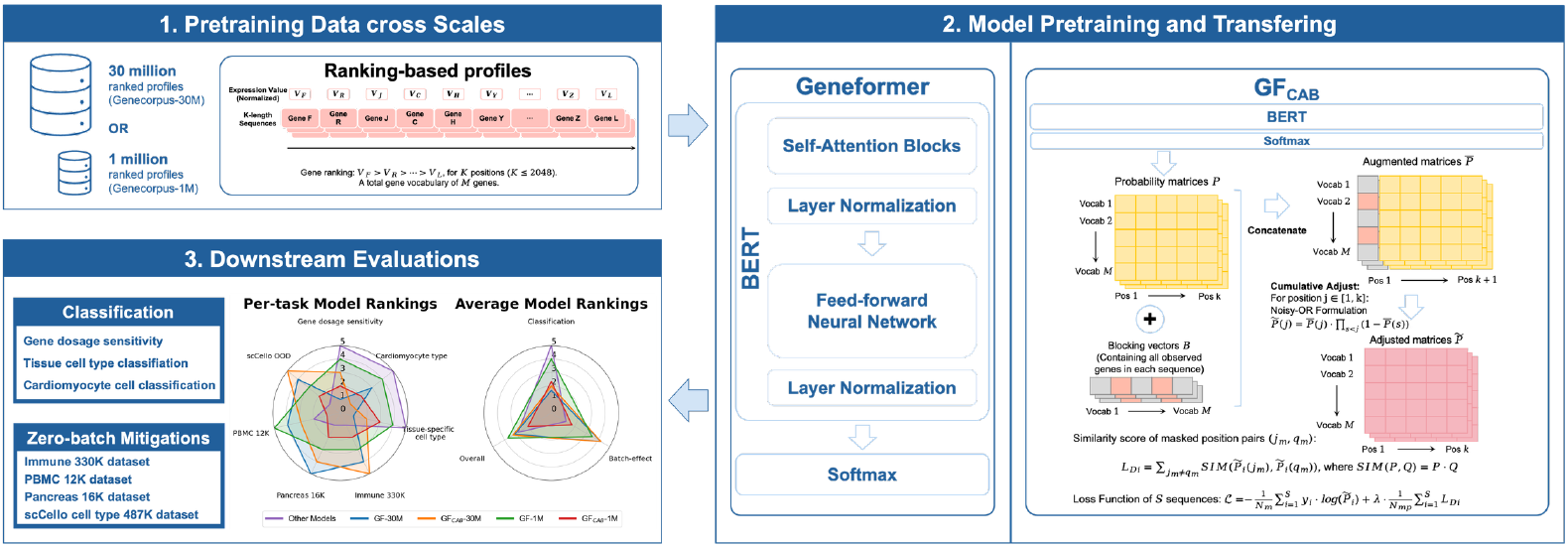
Study summary overview. Foundation models Geneformer (GF) and GF_CAB_ were pretrained on ranking-based single-cell transcriptomic profiles of different scales to evaluate the impact of data size. The models were thoroughly assessed on downstream classification tasks and zero-shot batch-effect mitigation tasks. Key results were further analyzed and compared to draw conclusions about the effect of pretraining scale and the suitability of ranking-based models for batch-effect mitigation.

**Fig 2.**
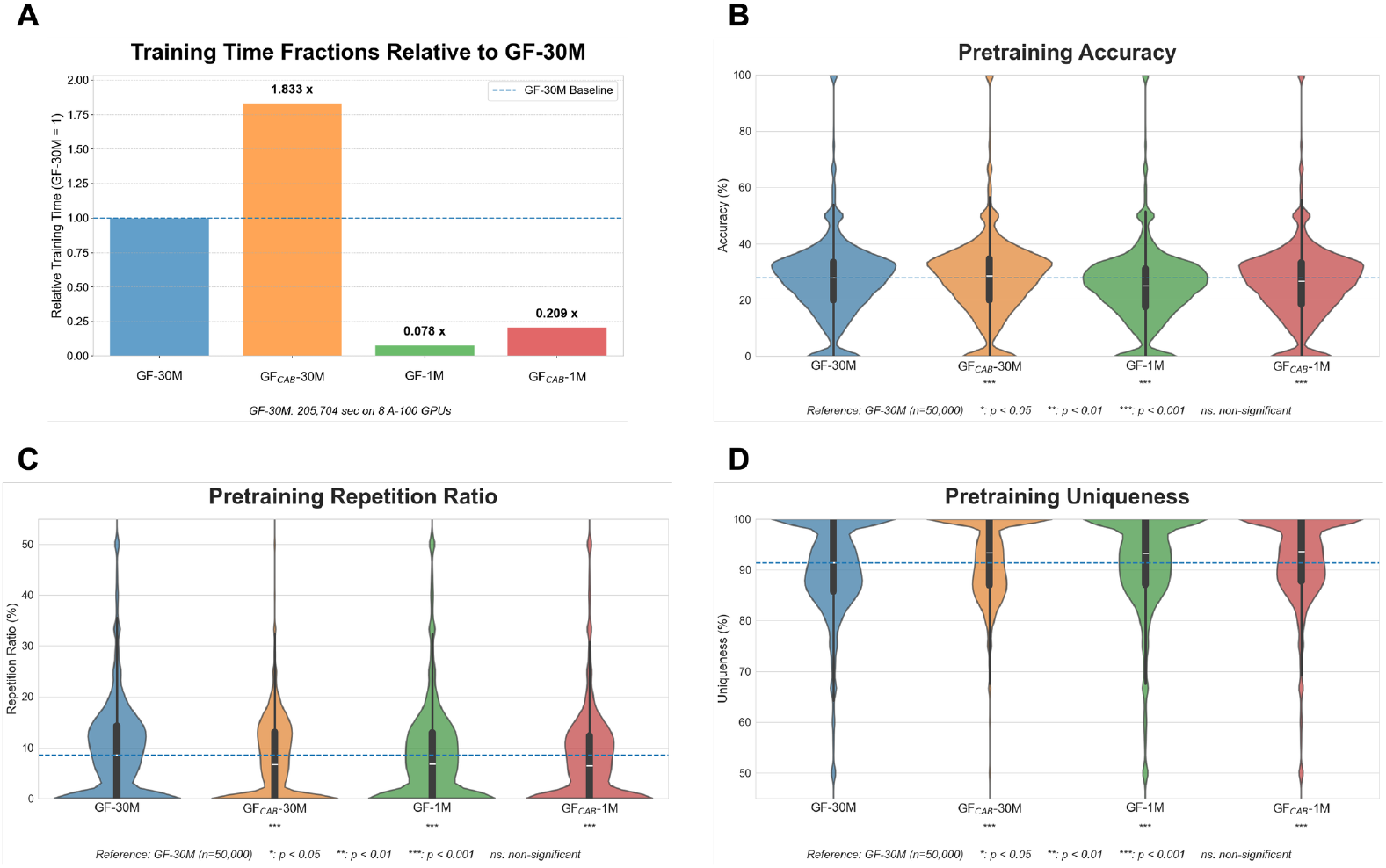
Relative training time and pretraining performance across models and data scales. (A) Models were pretrained on 8 A100 GPUs under the same settings, and relative training times were computed with respect to GF trained on 30M cells (GF-30M). (B) Distributions of pretraining accuracy across GF and GF_CAB_ models at the 30M and 1M data scales; Wilcoxon tests were performed against GF-30M, and white lines indicated the median of each violin plot. (C) Distributions of overall repetition ratios across GF and GF_CAB_ models at both scales; Wilcoxon tests were conducted against GF-30M. (D) Distributions of uniqueness scores across GF and GF_CAB_ models at both scales; Wilcoxon tests were conducted against GF-30M.

Across data scales, models trained on smaller corpora consistently produced more diverse predictions, with a higher proportion of distinct genes and reduced repetition among masked sites. This trend was observed in both GF and GF_CAB_, suggesting that scaling the pre-training dataset did not alleviate the tendency of rank-ordered models to recycle predictions from frequent tokens. Instead, smaller-scale models achieved greater exploratory diversity, uncovering less frequent but biologically important genes, while consuming only a fraction of computational resources. For example, models pre-trained on 1M profiles required approximately 7.8% and 20% of the resources relative to GF-30M, yet exhibited substantially higher prediction diversity and lower redundancy.

However, this efficiency and diversity came at the cost of reduced masked token prediction accuracy, reflecting a trade-off between scale and predictive fidelity. Crucially, evaluations on downstream tasks revealed that this loss of pre-training accuracy did not necessarily translate into inferior performance, underscoring the limited utility of accuracy as a sole indicator of generalization. Importantly, the GF_CAB_ variants compensated for much of the predictive loss at smaller scales: the GF_CAB_-1M model not only recovered masked token performance relative to its GF counterpart, but also achieved higher predictive diversity, highlighting the dual advantage of GF_CAB_ in improving prediction diversity and reducing reliance on large-scale pre-training, thus offering a more resource-efficient alternative for modeling single-cell transcriptomic data.

### Downstream performance

To assess generalization, we evaluated models pre-trained on both 30M and 1M profiles in terms of classification and zero-shot batch-effect tasks. Following Theodoris *et al*. [6], we tested three classification benchmarks, and according to Kedzierska *et al*. [17], we examined mitigation of batch effects in multiple datasets.

As shown in Fig 3, downstream outcomes varied by task type. For classification benchmarks, models pre-trained on the larger 30M corpus generally achieved stronger discriminative performance, reflecting the benefits of scale for fine-grained resolution of cell identities. In contrast, batch-effect evaluation favored models trained on smaller corpora, which exhibited superior robustness and generalization across heterogeneous datasets. This divergence confirmed that the scaling of pre-training data did not produce consistent improvements across biological tasks.

**Fig 3.**
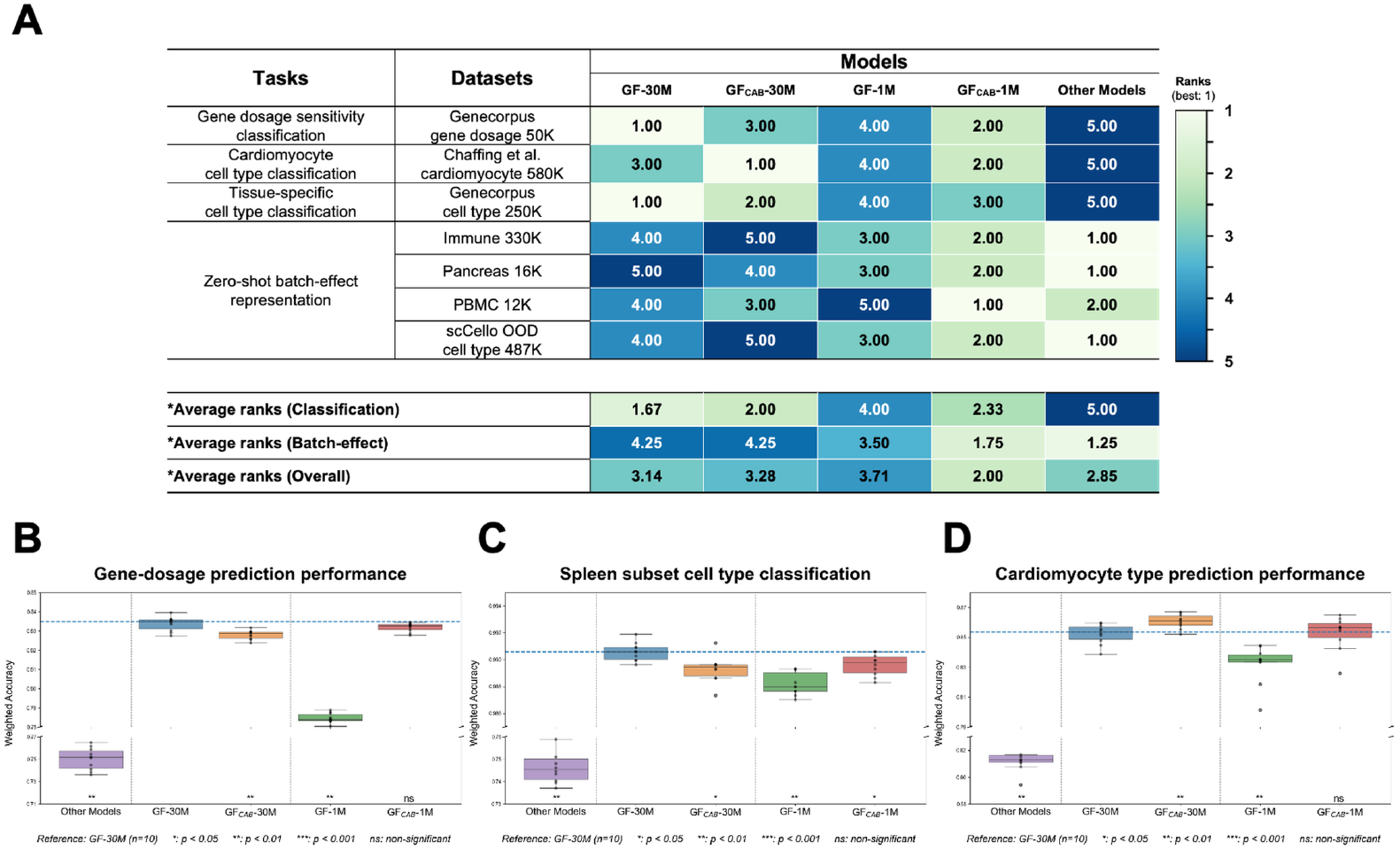
Downstream ranking results and classification performance across models. (A) Overall ranking results for GF and GF_CAB_ pretrained on 30M and 1M corpora, alongside baseline models, across three classification tasks and zero-shot batch-effect mitigation tasks on four datasets. (B) Gene dosage-sensitivity classification performance of GF, GF_CAB_, and baseline models was benchmarked, with Wilcoxon tests conducted relative to GF-30M. (C) Spleen cell type classification performance of GF, GF_CAB_, and baseline models was benchmarked, with Wilcoxon tests conducted relative to GF-30M. (D) Cardiomyocyte cell subtype classification performance of GF, GF_CAB_, and baseline models was benchmarked, with Wilcoxon tests conducted relative to GF-30M.

It should be noted that GF_CAB_ deviated from this trade-off. GF_CAB_-1M retained classification accuracy comparable to GF-30M while achieving markedly higher batch-effect mitigation performance. These results suggested that the structural refinements introduced in GF_CAB_ enabled better alignment of rank-ordered profiles and reduced dependence on the scale of the pre-training data, thus offering more balanced generalization across diverse applications.

### Classification performance

In classification benchmarks, models pre-trained on 30M profiles generally outperformed those trained on 1M profiles (Fig 3A). This pattern reflected the advantage of scale for discriminative tasks requiring fine-grained resolution of cell identity. Consistent with previous findings, task-specific baselines performed substantially worse than foundation models, underscoring the utility of large-scale pre-training for general-purpose classification.

A closer task-wise analysis (Fig 3B–D) revealed that this advantage did not hold uniformly across architectures. Although GF benefited noticeably from 30M compared to 1M, GF_CAB_-1M achieved performance comparable to GF-30M in two of the three benchmarks (detailed in S2 Table–S5 Table), effectively narrowing the expected gap between models trained on corpora differing by an order of magnitude. Furthermore, on the same scale, GF_CAB_-1M consistently surpassed GF-1M, showing that architectural refinements improved predictive alignment and reduced dependency on data scale.

These results indicated that careful architectural design could partially compensate for the lack of large-scale pre-training, challenging the assumption that classification gains primarily stem from data size.

### Zero-shot batch-effect mitigation performance

In zero-shot batch-effect evaluations (Fig 3A, S6 Table), scaling the pre-training corpus did not improve performance. In contrast, smaller-scale models consistently outperformed their larger counterparts in heterogeneous datasets, suggesting that reliance on large-scale pretraining reduced flexibility and impaired generalization across distinct cellular contexts for ranking-based foundation models. Moreover, compared to domain-specific methods such as scVI, both GF and GF_CAB_ performed substantially worse, highlighting the challenges of using rank-based representations for robust batch correction.

However, architectural refinements offered measurable improvements. GF_CAB_-1M outperformed GF across all tested datasets, narrowing the gap between the foundation models and the specialized, domain-specific methods. These gains reflected the improved alignment of GF_CAB_ with the rank-order profile structure, which reduced, but did not eliminate, the shortcomings of foundation models in this setting. Overall, these results indicated that the ranking of gene expression alone was insufficient for reliable batch correction and that future models may require representations explicitly designed with batch variability in mind.

Taken together, our downstream experiments revealed that larger pre-training corpora did not uniformly translate into stronger performance on various downstream tasks. Although classification tasks benefited from increased scale, batch-effect mitigation favored smaller models, and GF_CAB_ partially reconciled these opposing trends by reducing its dependence on the size of the data. Despite these improvements, rank-order foundation models continued to trail domain-specific bioinformatics tools in batch correction. These findings emphasized that architecture plays an equally critical role as data scale and that optimizing model design for biological consistency may be more impactful than further increasing the size of pre-training datasets.

### Ablation study

Next, we performed an ablation analysis to evaluate the contributions of cumulative prediction adjustment and similarity-based regularization in improving diversity in predictions. As summarized in Fig 4 and S7 Table, we compared GF with two modified variants: GF+PreAdj. (only with cumulative prediction adjustment) and GF_CAB_ (cumulative adjustment plus similarity-based regularization). Performance was assessed across pre-training, classification, and batch-effect mitigation, using models pre-trained on both 1M and 30M profiles.

**Fig 4.**
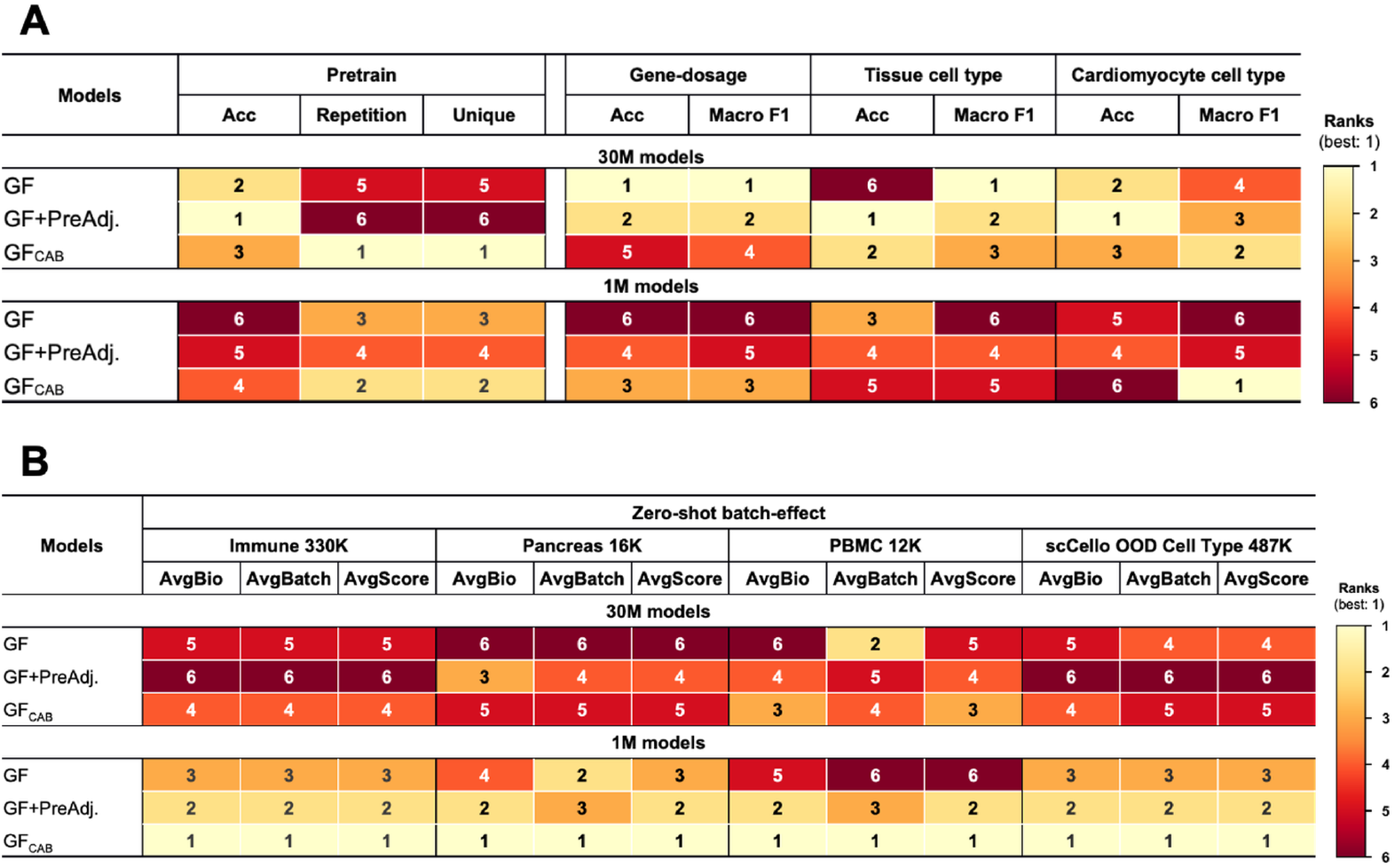
Ablation study results. GF and GF_CAB_ variants, GF+PreAdj. (with cumulative prediction adjustment only) and GF_CAB_ (with cumulative adjustment and similarity-based regularization loss), were pretrained on 30M and 1M datasets and evaluated on the pretraining objective, downstream classification, and batch-effect mitigation tasks. (A) Ranking results for pretraining performance and downstream classification tasks. (B) Ranking results for zero-shot batch-effect mitigation tasks.

For the pretraining objectives, introducing both cumulative adjustment and similarity-based loss yielded consistent improvements in accuracy and diversity compared to the GF on the same scale. In contrast, cumulative prediction adjustment alone showed weaker gains and even slightly underperformed GF at both scales, which is likely because the enlarged search space already increased the chance of correct predictions without additional adjustment. In comparison, the full GF_CAB_ architecture maintained clear benefits across scales, underscoring the complementary contributions of cumulative prediction adjustment and the similarity-based loss to improve pre-training. In classification tasks, both GF+PreAdj. and GF_CAB_ generally outperformed their GF counterparts across data scales, but improvements varied by task. Although the similarity-based penalty did not produce substantial accuracy gains beyond cumulative prediction adjustment, it preserved comparable classification performance while further enhancing prediction diversity (S7 Table). This suggests that the similarity loss acted as a stabilizing component, alleviating predictive redundancy without compromising discriminative ability.

The clearest benefits emerged in batch-effect evaluations. In all four data sets, both mechanisms progressively improved zero-shot generalization compared to GF, with GF_CAB_ delivering the strongest overall performance. These results confirmed that cumulative prediction adjustment and similarity-based regularization effectively alleviated the limitations of rank-based models in handling cross-dataset variability, thus mitigating the poor generalization noted in previous studies.

## Discussion

This work aimed to examine two fundamental questions in the development of foundation models for single-cell transcriptomics: (1) does scaling pre-training data consistently improve downstream performance, and (2) do models trained on rank-ordered profiles confer inherent advantages for batch effect mitigation? Our findings provide nuanced answers to both questions, challenging prevailing assumptions and pointing toward new design principles for biologically grounded model development.

With respect to the first question, scaling pretraining data improved performance in classification tasks for GF, echoing patterns observed in natural language processing, where larger corpora often yield finer-grained discriminative capacity. However, this scale advantage proved to be less robust than expected. The GF_CAB_ architecture, incorporating cumulative prediction adjustment and similarity-based regularization, achieved classification performance at the 1 million-profile scale that is comparable to or outperforms the GF pre-trained on 30M profiles. These results demonstrate that architectural alignment with the rank-ordered structure of gene expression profiles can offset or even replace the benefits of large-scale data. Thus, rather than assuming scale as the primary driver of downstream gains, our findings highlight the critical role of model design in ensuring the efficient use of biological information.

The second question revealed a more sobering outcome. Rank-based foundation models, including GF_CAB_, did not exhibit intrinsic advantages for zero-shot batch-effect mitigation. In fact, performance declined with larger pretraining corpora, underscoring a trade-off between scale-driven specialization and generalization across batches.

Although architectural refinements improved robustness compared to the baseline GF, these models continued to underperform relative to domain-specific methods such as scVI. These findings suggest that rank-based normalization alone is insufficient to capture the complex and context-dependent structure of batch effects. Dedicated objectives or representations explicitly tailored for cross-batch alignment may, therefore, be necessary to deliver state-of-the-art correction.

Together, these insights motivate a re-evaluation of the assumptions guiding the development of foundation models for single-cell data. First, more data is not always better: Scaling must be paired with architectural innovations that reflect the inherent structure of biological profiles. Second, rank-ordering strategies, while valuable for predictive diversity, are not a universal solution, particularly in settings where biological and technical variability are intertwined. Future research should pursue hybrid strategies that combine rank-based modeling with batch-sensitive objectives, explore multimodal integration to provide richer contextual signals, and investigate whether the observed interactions between scale and architecture generalize across other single-cell modalities and tissues.

In conclusion, our study underscores that the next generation of foundation models in single-cell biology will not be driven by scale alone. Instead, progress will hinge on architectures and objectives that faithfully represent the statistical and biological properties of single-cell data, thereby enabling models that are not only powerful, but also data-efficient and biologically robust.

## Methods

### Datasets

For full-scale pre-training, we followed the protocol established in Geneformer (GF) and used the Genecorpus-30M dataset [6], which contains 29,900,531 single-cell transcriptomic profiles, where the genes in each cell were ranked by normalized expression values. To examine the impact of reduced data scale, we constructed Genecorpus-1M by randomly sampling one million profiles from the original corpus. In addition, we curated a non-overlapping dataset of five million profiles (Genecorpus-5M), which was reserved exclusively for the unbiased evaluation of the masked pretraining objective.

Downstream benchmarking relied on datasets that were previously used in the GF study to ensure comparability across tasks. The gene dosage-sensitivity task was defined using 122 dosage-sensitive and 368 dosage-insensitive genes reported in previous studies [21–23], along with 50,000 ranked profiles curated by Theodoris et al. [6] (Genecorpus gene dosage 50K). Multi-tissue cell type classification was performed on 249,556 profiles spanning 59 cell types across nine tissues (spleen, kidney, lung, liver, brain, placenta, large intestine, immune system, and pancreas) [6] (Genecorpus cell type 250K). For cardiomyocyte disease-state classification, we used the profiles from Chaffin et al. [24], which included cardiomyocytes from non-failing hearts, as well as from patients with hypertrophic cardiomyopathy (HCM) [25] and dilated cardiomyopathy (DCM) [26]. This dataset encompassed 580,000 ranked profiles from 42 individuals, annotated with metadata including age, sex, and cell type, producing a three-class classification task (Chaffin cardiomyocyte 580K).

To evaluate the quality of learned cell embeddings and assess batch integration under zero-shot conditions, we incorporated four external datasets following prior work [15, 17]: Pancreas 16K [27], PBMC 12K [28, 29], Immune 330K [30], and scCello OOD cell type 487K [15].

### Enhancing prediction diversity with GF_**CAB**_

Geneformer pioneered the application of language models (LMs) for single-cell RNA-seq by representing gene co-expression patterns as rank-ordered sequences [6]. Although effective, its masked language modeling (MLM) framework exhibits three limitations: (1) repeated assignment of the same gene across multiple masked positions within a profile, (2) systematic under-representation of low-expression genes, and (3) frequency-driven bias toward highly expressed, ubiquitous genes. To address these issues, we propose **GF**_**CAB**_ (Geneformer with **C**umulative-assignment on the prediction **A**djustment and **B**oosting on the similarity-based diversity), which augments GF with two complementary mechanisms: cumulative-assignment prediction adjustment and similarity-based regularization.

### Cumulative-assignment prediction adjustment

In standard MLM, masked positions are predicted independently, allowing the same gene to be assigned repeatedly across multiple sites. To mitigate this redundancy, we introduce a cumulative assignment-and-blocking mechanism that integrates one-hot substitution with cumulative Noisy-OR suppression [31].

Given a masked sequence 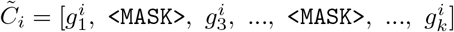 of *k* tokens, the BERT model produces a *k × V* probability matrix *P*_*i*_, where *V* ∈ ℝ is the vocabulary size. Observed positions are replaced with one-hot vectors: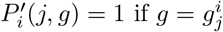, and 0 otherwise. A blocking vector *B*_*i*_ ∈ {0, 1}^*V*^ is then defined such that *B*_*i*_(*g*) = 1 if *g* ∈ *G*_observed_ (observed gene set within a profile), and it is concatenated with 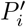 to form the augmented matrix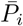, as shown in Eq (1).

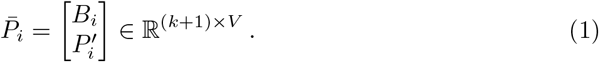

After complement 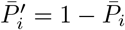, Noisy-OR normalization produces adjusted probabilities 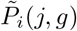, as shown in Eq (2), which enforces cumulative suppression across positions. This mechanism mitigates duplicate predictions within the same cell profile, thereby improving biological plausibility.

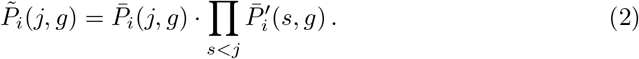

### Similarity-based regularization

Despite the redundancy observed within individual cell profiles, the predictions made by the GF remain strongly biased toward highly expressed, ubiquitous genes. This pattern suggests an overreliance on global expression trends, with limited utilization of local, context-specific signals that are essential for meaningful biological interpretation.

To address this frequency bias limitation, we introduce a similarity-based regularization strategy, inspired by Ayinde *et al*. [32], which is designed to penalize overly similar predictions across masked positions and encourage context-aware inference. This approach directly counteracts the model’s tendency to favor frequent and highly expressed genes, thus improving its ability to capture rare, condition-specific signals and consequently improving predictive diversity.

For masked positions (*j*_*m*_, *q*_*m*_) in sequence 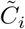, we compute pairwise similarity between adjusted distributions, as shown in Eq (3).

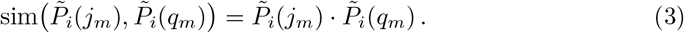

Summing across all *N*_*mp*_ distinct masked-pair combinations produces a diversity loss, Eq (4), which discourages uniform predictions across masked sites by penalizing distributional redundancy.

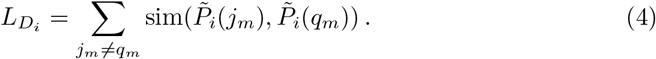

Finally, we incorporate this similarity penalty into the masked sequence pretraining objective Eq (5) by combining it with the standard cross-entropy loss. Here, *S* ∈ ℝ, *N*_*m*_ ∈ ℝ, and *y*_*i*_ ∈ ℝ^*V*^ denote the number of sequences, the number of masked positions, and the one-hot representation of the ground-truth label for sequence *i*, respectively.

The hyperparameter *λ* ∈ ℝ controls the strength of diversity regularization. This penalty discourages over-concentration on frequent genes, promoting rare and context-sensitive predictions that are critical for biological discovery.

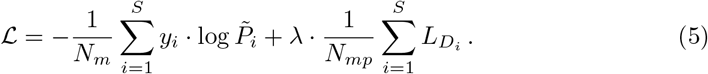

### Baseline models

We compared GF_CAB_ against GF in three dimensions: pre-training objectives, downstream classification, and zero-shot batch-effect mitigation. In classification tasks, we also included Support Vector Machine (SVM) [33], Random Forest (RF) [34], and Logistic Regression (LR) as baselines. For zero-shot settings, GF and GF_CAB_ were compared with highly variable genes (HVG) and scVI [35], a variational autoencoder framework that explicitly models batch effects.

### Evaluation metrics

For pretraining evaluation, models were evaluated on the masked gene prediction objective using metrics that jointly capture predictive accuracy and diversity. Masked prediction accuracy quantifies the proportion of correctly reconstructed masked tokens across the validation set, reflecting the model’s ability to recover contextually appropriate genes. Repetition ratios measure redundancy in predictions and are reported in three forms: the overall repetition ratio (fraction of all duplicate predictions), the unmasked repetition ratio (proportion of predictions overlapping with unmasked input tokens), and the masked repetition ratio (fraction of repeated predictions restricted to masked positions). To further characterize predictive diversity, we introduced a uniqueness score, inspired by entropy-based measures such as the Kullback–Leibler divergence [36]. This score reflects the proportion of distinct gene predictions at masked sites, capturing the model’s ability to avoid mode collapse and explore less frequent, yet biologically informative genes. Higher uniqueness values indicate more diverse and context-sensitive predictions.

For downstream tasks, we used classification, clustering, and batch integration metrics to jointly assess biological relevance and technical robustness. Classification performance was evaluated by accuracy, macro-averaged AUC [37, 38], macro and weighted F1 scores [39, 40], and recall [41], complemented by confusion matrices for error-structure visualization [42]. Cluster quality, following established benchmarking frameworks [17, 27], was quantified using normalized mutual information (NMI) [43], adjusted Rand index (ARI) [44], and average silhouette width (ASW_label_), which were aggregated into the AvgBIO metric [8] to summarize cluster separability. Batch integration was assessed through the average silhouette width with respect to batch (ASW_batch_) [45] and graph connectivity (GraphCon) [15], which were combined into an AvgBatch measure. To provide a balanced evaluation, we reported an overall score (AvgScore) [8], defined as the mean of AvgBIO and AvgBatch, which integrate biological fidelity with batch mixing quality.

### Implementation details

We implemented the proposed architecture, GF_CAB_, as an extension of the masked language model GF. The model retains GF’s core transformer design (6 encoder layers, 4 attention heads, 256-dimensional embeddings, 512-dimensional feedforward layers, and a maximum input length of 2,048) while introducing cumulative prediction adjustment and a similarity-based loss penalty for diversity boosting. Hyperparameters, training schedules, and preprocessing strategies were aligned with GF to enable direct comparison (see S8 Table). Input preparation followed the standard masking scheme through a modified Hugging Face Trainer framework [46], with 15% of tokens selected for perturbation and custom batching to account for variable gene counts per cell. Pre-training was conducted at two scales: a 1M-cell dataset (Genecorpus-1M) with a batch size of 12 for rapid iteration and a 30M-cell dataset (Genecorpus-30M) with a batch size of 8 for full-scale training, both run on 8 NVIDIA A100 GPUs. GF was pretrained under identical conditions to ensure a controlled baseline.

For downstream evaluation, we fine-tuned GF, GF_CAB_, and classical machine learning baselines (SVM, random forest, and logistic regression) on three classification tasks that reflect distinct biological challenges. Depending on task requirements, we used either Hugging Face’s `BertForTokenClassification` for gene-level prediction or `BertForSequenceClassification` for cell profile–level tasks, initializing with pre-trained weights and applying GF’s fine-tuning conventions. Hyperparameters such as frozen layers, learning rate schedules, warmup steps, and epochs were set according to the GF’s released scripts, with minor task-specific adjustments (*e*.*g*., different frozen layer numbers and learning rates for cardiomyocyte classification). Baseline classifiers were implemented using scikit-learn [47] with default parameters, except for training epochs, which were harmonized with task objectives.

To test zero-shot transferability, we compared the embeddings from GF_CAB_ and GF with those derived from highly variable genes (HVG) and scVI, following the protocol established by Kedzierska *et al*. [17]. For the transformer models, cell embeddings were computed from unmasked gene inputs without further masking or fine-tuning, ensuring strict zero-shot conditions. HVG features provided a conventional dimensionality reduction baseline, while scVI embeddings were drawn directly from the latent space. All embeddings were benchmarked using cluster- and integration-based metrics, allowing for a direct comparison between foundation models and established single-cell representation methods.

### Statistical analysis

Pre-training results are reported as the mean *±* standard deviation across 50,000 independent ranked single-cell profiles. For downstream tasks, accuracy and macro-F1 values were averaged over 10 independent cross-validation runs (where applicable). Statistical significance of performance differences was evaluated using the paired Wilcoxon signed-rank test, with each model compared pairwise against GF-30M as the baseline, unless otherwise noted.

## Reproducibility

All experiments were performed using PyTorch (v2.6.0) with Hugging Face Transformers (v4.38.2), CUDA (v11.8), and Python (v3.11.7). Training was distributed across 8 NVIDIA A100 GPUs with mixed precision enabled. Random seeds were fixed across NumPy, PyTorch, and CUDA to ensure replicability. The complete codebase, pretrained weights, evaluation scripts, and datasets incorporated in this study are available at the following repositories.

## Data Availability

All datasets used in this study are publicly available in the Hugging Face repository at https://huggingface.co/datasets/JFLa/GF-CAB_Datasets. This includes the pretraining datasets at two scales, the evaluation datasets, the downstream classification datasets, and the zero-shot batch-effect correction evaluation datasets.

## Code Availability

The code scripts used for model training, evaluation, and analysis are available in both the GitHub repository at https://github.com/CHENJ-VV/GF-CAB.git and the Hugging Face repository at https://huggingface.co/JFLa/GF-CAB. The repository includes scripts for data processing, model training, and benchmarking results generation.

## Acknowledgments

We would like to acknowledge Xin Gao and Jesper Tegnér for their valuable feedback and constructive discussions that helped strengthen this study. We would also like to thank the Department of Biostatistics and Bioinformatics at Duke University for their collaborative insights and the KAUST Supercomputing Core Laboratory for providing essential computational resources used in model pretraining and large-scale analysis.

## Funding

This work was supported by the Smart Health Initiative and baseline-funding from the King Abdullah University of Science and Technology (KAUST).

## Supporting information

**S1 Table. Pretraining performance of models on 50**,**000 profiles from Genecorpus-5M**. Pretraining results are reported for Geneformer and GF_CAB_ models with varying similarity regularization coefficients (*λ*), trained on different scales of pretraining data and evaluated on 50,000 test profiles randomly sampled from Genecorpus-5M. Metrics include masked prediction accuracy (mean *±* standard deviation), repetition ratios (overall, unmasked, and masked), and uniqueness scores.

**S2 Table. Gene dosage-sensitivity prediction performance of Geneformer and GF**_**CAB**_ **models**. Macro F1-scores and accuracies are reported for Geneformer and GF_CAB_ variants with different similarity regularization coefficients (*λ*), pretrained on either Genecorpus-30M or Genecorpus-1M. Bold values indicate the best performance, while underlined values indicate the second-best performance.

**S3 Table. Cardiomyocyte cell type prediction performance of Geneformer and GF**_**CAB**_ **models**. Macro F1-scores and accuracies are reported for Geneformer and GF_CAB_ variants with different similarity regularization coefficients (*λ*), pretrained on either Genecorpus-30M or Genecorpus-1M. Bold values indicate the best performance, while underlined values indicate the second-best performance.

**S4 Table. Tissue-specific cell type prediction performance of Geneformer and GF**_**CAB**_ **models**. Macro F1-scores and accuracies are reported for individual tissues using Geneformer and GF_CAB_ variants with different similarity regularization coefficients (*λ*), pretrained on either Genecorpus-30M or Genecorpus-1M. Bold values denote the best performance, while underlined values denote the second-best performance.

**S5 Table. Spleen-specific cell type prediction performance of Geneformer and GF**_**CAB**_ **models**. Macro F1-scores and accuracies are reported for spleen-derived cells using Geneformer and GF_CAB_ variants with different similarity regularization coefficients (*λ*), pretrained on either Genecorpus-30M or Genecorpus-1M. Bold values denote the best performance, while underlined values denote the second-best performance.

**S6 Table. Zero-shot batch-effect mitigation performance of Geneformer and GF**_**CAB**_ **models**. Batch-effect mitigation performance is reported for Geneformer and GF_CAB_ variants with different similarity regularization coefficients (*λ*), pretrained on either Genecorpus-30M or Genecorpus-1M. Evaluation metrics include cell type clustering measures (NMI, ARI, ASW_label_, AvgBio), batch clustering measures (Graph connectivity score, ASW_batch_, AvgBatch), and the overall score (average of AvgBio and AvgBatch).

**S7 Table. Ablation study results for Geneformer and GF**_**CAB**_ **models**. Ablation results are reported for Geneformer and GF_CAB_ variants with different similarity regularization coefficients (*λ*), pretrained on either Genecorpus-30M or Genecorpus-1M. Evaluations span pretraining performance, gene dosage-sensitivity prediction, tissue-specific and cardiomyocyte cell type classification, and zero-shot batch-effect mitigation across four datasets using the metrics described above. Bold values denote the best performance, while underlined values denote the second-best performance.

**S8 Table. Pretraining hyperparameters for Geneformer and GF**_**CAB**_ **models**. Detailed hyperparameter configurations used for Geneformer and GF_CAB_ models benchmarked in this study are reported.

## Notes

### Competing Interest Statement

The authors have declared no competing interest.

## References

1. Bommasani R, Hudson DA, Adeli E, Altman R, Arora S, von Arx S, et al. On the Opportunities and Risks of Foundation Models [Preprint]. arXiv; 2021. Available from: https://arxiv.org/abs/2108.07258.

2. Brown TB, Mann B, Ryder N, Subbiah M, Kaplan J, Dhariwal P, et al. Language Models are Few-Shot Learners [Preprint]. arXiv; 2020. Available from: https://arxiv.org/abs/2005.14165.

3. Ramesh A, Pavlov M, Goh G, Gray S, Voss C, Radford A, et al. Zero-Shot Text-to-Image Generation [Preprint]. arXiv; 2021. Available from: https://arxiv.org/abs/2102.12092.

4. Tang F, Barbacioru C, Wang Y, Nordman E, Lee C, Xu N, et al. mRNA-Seq whole-transcriptome analysis of a single cell. Nat Meth. 2009 Apr;6:377–82. doi:10.1038/nmeth.1315.

5. Navin N, Kendall J, Troge J, Andrews P, Rodgers L, McIndoo J, et al. Tumour evolution inferred by single-cell sequencing. Nature. 2011 Mar;472:90–4. doi:10.1038/nature09807.

6. Theodoris CV, Xiao L, Chopra A, Chaffin MD, Al Sayed ZR, Hill MC, et al. Transfer learning enables predictions in network biology. Nature. 2023 May;618:616–24. doi:10.1038/s41586-023-06139-9.

7. Yang F, Wang W, Wang F, Fang Y, Tang D, Huang J, et al. scBERT as a large-scale pretrained deep language model for cell type annotation of single-cell RNA-seq data. Nat Mach Intell. 2022 Sep;4:852–66. doi:10.1038/s42256-022-00534-z.

8. Cui H, Wang C, Maan H, Pang K, Luo F, Duan N, et al. scGPT: toward building a foundation model for single-cell multi-omics using generative AI. Nat Meth. 2024 Feb;21:1470–80. doi:10.1038/s41592-024-02201-0.

9. Hao M, Gong J, Zeng X, Liu C, Guo Y, Cheng X, et al. Large-scale foundation model on single-cell transcriptomics. Nat Meth. 2024 Jun;21:1481–91. doi:10.1038/s41592-024-02305-7.

10. Devlin J, Chang MW, Lee K, Toutanova K. BERT: Pre-training of Deep Bidirectional Transformers for Language Understanding [Preprint]. arXiv; 2018. Available from: https://arxiv.org/abs/1810.04805.

11. Roohani Y, Huang K, Leskovec J. Predicting transcriptional outcomes of novel multigene perturbations with GEARS. Nat Biotechnol. 2024 Jun;42(6):927–35. doi:10.1038/s41587-023-01905-6.

12. Wenteler A, Occhetta M, Branson N, Huebner M, et al. PertEval-scFM: Benchmarking Single-Cell Foundation Models for Perturbation Effect Prediction [Preprint]. bioRxiv; 2024. Available from: https://www.biorxiv.org/content/10.1101/2024.10.02.616248v1.

13. Soneson C, Robinson MD. Bias, robustness and scalability in single-cell differential expression analysis. Nat Meth. 2018 Apr;15(4):255–61. doi:10.1038/nmeth.4612.

14. Xu C, Lopez R, Mehlman E, et al. Probabilistic harmonization and annotation of single-cell transcriptomics data with deep generative models. Mol Syst Biol. 2021;17(1):e9620. doi:10.15252/msb.20209620.

15. Yuan X, Zhan Z, Zhang Z, Zhou M, Zhao J, Han B, et al. Cell-ontology guided transcriptome foundation model [Preprint]. arXiv; 2024. Available from: https://arxiv.org/abs/2408.12373.

16. Kedzierska KZ, Crawford L, Amini AP, et al. Assessing the limits of zero-shot foundation models in single-cell biology [Preprint]. bioRxiv; 2023. Available from: https://www.biorxiv.org/content/10.1101/2023.10.16.561085v1.

17. Kedzierska KZ, Crawford L, Amini AP, et al. Zero-shot evaluation reveals limitations of single-cell foundation models. Genome Biol. 2025 Apr;26(1):101. doi:10.1186/s13059-025-03574-x.

18. Mitchell J, Goyal S, Wen K, et al. Overtrained Language Models Are Harder to Fine-Tune [Preprint]. arXiv; 2025. Available from: https://arxiv.org/abs/2503.19206.

19. Mizrahi D, Lindbo Larsen AB, Allardice J, Petryk S, Gorokhov Y, Li J, et al. Language Models Improve When Pretraining Data Matches Target Tasks [Preprint]. arXiv; 2025. Available from: https://arxiv.org/abs/2507.12466.

20. Chen Z, Wang S, Xiao T, Wang Y, Chen S, Cai X, et al. Revisiting Scaling Laws for Language Models: The Role of Data Quality and Training Strategies. In: Che W, Nabende J, Shutova E, Pilehvar MT, editors. Proceedings of the 63rd Annual Meeting of the Association for Computational Linguistics (Volume 1: Long Papers). Vienna, Austria: Association for Computational Linguistics; 2025. p. 23881–99. doi:10.18653/v1/2025.acl-long.1163.

21. Exome Aggregation Consortium, Lek M, Karczewski KJ, Minikel EV, Samocha KE, Banks E, et al. Analysis of protein-coding genetic variation in 60,706 humans. Nature. 2016 Aug;536(7616):285–91. Available from: https://www.nature.com/articles/nature19057. doi:10.1038/nature19057.

22. Shihab HA, Rogers MF, Campbell C, Gaunt TR. HIPred: an integrative approach to predicting haploinsufficient genes. Bioinformatics. 2017 Jun;33(12):1751–7. doi:10.1093/bioinformatics/btx028.

23. Ni Z, Zhou XY, Aslam S, Niu DK. Characterization of Human Dosage-Sensitive Transcription Factor Genes. Frontiers in Genetics. 2019. doi:10.3389/fgene.2019.01208.

24. Chaffin M, Papangeli I, Simonson B, Akkad AD, Hill MC, Arduini A, et al. Single-nucleus profiling of human dilated and hypertrophic cardiomyopathy. Nature. 2022 Aug;608(7921):174–80. Available from: https://www.nature.com/articles/s41586-022-04817-8. doi:10.1038/s41586-022-04817-8.

25. Liu X, Ma Y, Yin K, Li W, Chen W, Zhang Y, et al. Long non-coding and coding RNA profiling using strand-specific RNA-seq in human hypertrophic cardiomyopathy. Scientific Data. 2019 Jun;6(1):90. Available from: https://www.nature.com/articles/s41597-019-0094-6. doi:10.1038/s41597-019-0094-6.

26. Sweet ME, Cocciolo A, Slavov D, Jones KL, Sweet JR, Graw SL, et al. Transcriptome analysis of human heart failure reveals dysregulated cell adhesion in dilated cardiomyopathy and activated immune pathways in ischemic heart failure. BMC Genomics. 2018 Nov;19(1):812. doi:10.1186/s12864-018-5213-9.

27. Luecken MD, Büttner M, Chaichoompu K, Danese A, … Theis FJ. Benchmarking atlas-level data integration in single-cell genomics. Nat Methods. 2022;19(1):41–50. doi:10.1038/s41592-021-01336-8.

28. Gayoso A, Lopez R, Xing G, Boyeau P, Amiri VVP, Hong J, et al. A Python library for probabilistic analysis of single-cell omics data. Nat Biotechnol. 2022;40(2):163–6. doi:10.1038/s41587-021-01206-w.

29. Zheng GXY, Terry JM, Belgrader P, Ryvkin P, Bent ZW, Wilson R, et al. Massively parallel digital transcriptional profiling of single cells. Nat Commun. 2017;8:14049. doi:10.1038/ncomms14049.

30. Domínguez-Conde C, Xu C, Jarvis LB, Rainbow DB, Wells SB, Gomes T, et al. Cross-tissue immune cell analysis reveals tissue-specific features in humans. Science. 2022 May;376(6594):eabl5197. doi:10.1126/science.abl5197.

31. Praveen P, Fröhlich H. Boosting Probabilistic Graphical Model Inference by Incorporating Prior Knowledge from Multiple Sources. PLOS ONE. 2013;8(6):e67410. doi:10.1371/journal.pone.0067410.

32. Ayinde BO, Inanc T, Zurada JM. Regularizing Deep Neural Networks by Enhancing Diversity in Feature Extraction. IEEE Transactions on Neural Networks and Learning Systems. 2019;30(9):2650–61. Available from: https://ieeexplore.ieee.org/document/8354782. doi:10.1109/TNNLS.2018.2878873.

33. Boser BE, Guyon IM, Vapnik VN. A Training Algorithm for Optimal Margin Classifiers. In: Proceedings of the 5th Annual Workshop on Computational Learning Theory (COLT ‘92). Pittsburgh, PA, USA: ACM Press; 1992. p. 144–52. doi:10.1145/130385.130401.

34. Breiman L. Random Forests. Machine Learning. 2001;45(1):5–32. doi:10.1023/A:1010933404324.

35. Lopez R, Regier J, Cole MB, Jordan MI, Yosef N. Deep generative modeling for single-cell transcriptomics. Nat Methods. 2018 Dec;15(12):1053–8. doi:10.1038/s41592-018-0229-2.

36. Kullback S, Leibler RA. On Information and Sufficiency. Annals of Mathematical Statistics. 1951;22(1):79–86. doi:10.1214/aoms/1177729694.

37. Hanley JA, McNeil BJ. The meaning and use of the area under a receiver operating characteristic (ROC) curve. Radiology. 1982 Apr;143(1):29–36. doi:10.1148/radiology.143.1.7063747.

38. Junge MRJ, Dettori JR. ROC Solid: Receiver Operator Characteristic (ROC) Curves as a Foundation for Better Diagnostic Tests. Global Spine J. 2018 Jun;8(4):424–9. doi:10.1177/2192568218778294.

39. Opitz J, Burst S. Macro F1 and Macro F1 [Preprint]. arXiv; 2019. Available from: https://arxiv.org/abs/1911.03347.

40. Taha AA, Hanbury A. Metrics for evaluating 3D medical image segmentation: analysis, selection, and tool. BMC Med Imaging. 2015 Aug;15:29. doi:10.1186/s12880-015-0068-x.

41. Saito T, Rehmsmeier M. The Precision-Recall Plot Is More Informative than the ROC Plot When Evaluating Binary Classifiers on Imbalanced Datasets. PLOS ONE. 2015;10(3):e0118432. doi:10.1371/journal.pone.0118432.

42. Stehman SV. Selecting and Interpreting Measures of Thematic Classification Accuracy. Remote Sensing of Environment. 1997;62:77–89. doi:10.1016/S0034-4257(97)00083-7.

43. Vinh NX, Epps J, Bailey J. Information Theoretic Measures for Clusterings Comparison: Is a Correction for Chance Necessary? In: Proceedings of the 26th International Conference on Machine Learning (ICML ‘09); 2009. p. 1073-80. doi:10.1145/1553374.1553511.

44. Hubert L, Arabie P. Comparing Partitions. Journal of Classification. 1985;2(1):193–218. doi:10.1007/BF01908075.

45. Tran HTN, Ang KS, Chevrier M, Zhang X, Lee NYS, Goh M, et al. A benchmark of batch-effect correction methods for single-cell RNA sequencing data. Genome Biology. 2020;21(1):12. doi:10.1186/s13059-019-1850-9.

46. Wolf T, Debut L, Sanh V, Chaumond J, Delangue C, Moi A, et al. HuggingFace’s Transformers: State-of-the-art Natural Language Processing. arXiv; 2020. Available from: https://arxiv.org/abs/1910.03771. 1910.03771.

47. Buitinck L, Louppe G, Blondel M, Pedregosa F, Mueller A, Grisel O, et al. API design for machine learning software: experiences from the scikit-learn project. arXiv preprint arXiv:13090238. 2013. Version 1. Available from: https://arxiv.org/abs/1309.0238. doi:10.48550/ARXIV.1309.0238.

